# Functional network dynamics in a neurodevelopmental disorder of known genetic origin

**DOI:** 10.1101/463323

**Authors:** Erin Hawkins, Danyal Akarca, Mengya Zhang, Diandra Brkic, Mark Woolrich, Kate Baker, Duncan Astle

**Affiliations:** MRC Cognition and Brain Sciences Unit, University of Cambridge, 15 Chaucer Road, Cambridge, CB2 7EF, UK; Oxford Centre for Human Brain Activity, University of Oxford, University Department of Psychiatry, Warneford Hospital, Oxford, OX3 7JX, UK; Department of Medical Genetics, University of Cambridge, Cambridge Institute for Medical Research, Cambridge, CB2 0XY, UK

**Keywords:** Atypical brain development, cognitive development, functional connectivity, human genetics, magnetoencephalography (MEG)

## Abstract

Cognitive processing depends on the temporal co-ordination of functional brain networks. This fundamental aspect of neurophysiology potentially bridges the genetic regulation of neuronal activity and developmental cognitive impairments. We investigated brain network dynamics in a neurodevelopmental disorder of known genetic origin, by comparing individuals with *ZDHHC9*-associated intellectual disability to individuals with no known impairment. We used Hidden Markov Modelling on magnetoencephalography (MEG) data, at rest and during auditory oddball stimulation, to characterise transient network dynamics. At rest, network dynamics distinguished the groups, with *ZDHHC9* participants showing longer state activation. Crucially, *ZDHHC9* gene expression levels predicted the group differences across networks, supporting an association between molecular pathology and neurophysiology. In contrast, network dynamics during auditory oddball stimulation did not show this association. We demonstrate a link between brain network dynamics and regional gene expression, and present a valuable method for understanding the real-time neural mechanisms linking genetic variation to cognitive difficulties.

## Introduction

In recent years, whole-brain imaging methods have advanced our ability to characterise functional brain connectivity, and have revealed that distributed neural systems support cognition and behaviour (Astle et al., 2016; Barnes et al., 2015; Smith et al., 2015; Vidaurre et al. 2017a). Disruptions to functional connectivity are considered a characteristic feature of multiple developmental disorders, but surprisingly little is known about the mechanisms that drive this variability. It is well known that genetic factors influence the intrinsic organisation of functional networks (e.g. Bathelt et al., 2016, 2017; Colclough et al., 2017; Wang et al., 2015). In many cases, genes are likely to influence the developmental emergence of functional networks via regulation of continuous activity-dependent physiological processes, rather than by fixed anatomical differences. However, understanding of the rapid synchronization and integration of functional connectivity networks on fast time-scales is limited, partly because there is scarcity of methods capable of characterising the fast dynamics of brain networks. As a result, there is currently little understanding of the genetic effects on these networks and the cellular mechanisms that could drive their developmental variability, or the consequences of perturbations to these network dynamics for cognition. The aim of this study is to redress this by exploring dynamic transient brain connectivity in a group of individuals with a neurodevelopmental disorder of known genetic origin.

The current methods capable of deriving system wide neural networks largely assume that the networks remain stationary over time. Functional magnetic resonance imaging (fMRI) has been the primary method for investigating coordination between brain regions, by measuring covariations in the haemodynamic response (Beckmann et al., 2005; Damoiseaux et al., 2006; Smith et al., 2009). Magnetoencephalography (MEG) provides a powerful complement to fMRI because it captures the electrophysiological nature of the underlying networks (Barnes et al. 2016). The primary approach to identifying functional connectivity networks with MEG is to measure the synchronisation between the amplitude envelopes of oscillations within certain frequency bands, which yields networks of co-ordinated activity across spatially separate brain regions (de Pasquale et al., 2010; O’Neill et al., 2015; Sudre et al., 2017). This approach has characterised resting state networks (RSNs) that show a closely overlapping topography with those found using fMRI (Astle et al., 2016; Barnes et al., 2015; Demuru et al., 2017; de Pasquale et al., 2010). However, with both fMRI and MEG, the estimates of functional connectivity are typically derived by aggregating activity over large time windows, such as the full resting state acquisition period. This defines large-scale networks as remaining stationary, whether during rest or task performance. If large-scale neural networks play an important role in supporting cognitive processes, then they must synchronise their activity on a timescale of around 50-200ms, switching far more quickly than can be captured using averages taken from time window methods (Hindriks et al., 2017).

More recently, researchers have developed analytical techniques that fully capitalise on the high temporal resolution of MEG. An example is the use of Hidden Markov Modelling (HMM) to explore the temporal dynamics of these networks (Baker et al., 2014; Vidaurre et al., 2016; Viduarre et al., 2017b; Vidaurre et al. in press). Instead of estimating the correlation between band-limited amplitude envelopes separately in one large time window at a time, the HMM is a data-driven method that identifies a sequence of “states”, whereby each state corresponds to a unique pattern of brain network activity and correlation that reoccurs at different points in time. For example, the pattern of amplitude envelope activity that characterises each state can be well-estimated by pooling over the evidence provided by the many repeated visits to the state. This means that individual state visits can be potentially very short in time. Indeed, it has been shown that the HMM can identify network dynamics in resting MEG on ~100ms time-scale, which is much faster than can be investigated with traditional sliding time window approaches (Baker et al., 2014; Viduarre et al., 2016). By quantifying the time-series of MEG data as a sequence of transient states, the HMM provides information about the points in time at which each state is active, enabling the characterisation of the temporal dynamics of each state or network. Further, the HMM allows the detection of rapid, transient organisation of brain networks and therefore represents a powerful tool for understanding how network dynamics can be influenced by, for example, genetic factors.

The application of these methods remains in its infancy. As a result we have yet to establish how underlying neurobiological or molecular mechanisms can influence the formation of these large-scale networks or their intrinsic temporal dynamics. Individuals with rare single gene mutations can represent a unique and interesting window to investigate the specific interactions between cellular and physiological processes involved in cognitive development. This is because, unlike neurodevelopmental disorders defined by behavioural impairments, they have a highly specific and known aetiology, common across all cases. Here, we studied a group of individuals with mutations in *ZDHHC9*, a rare recurrent cause of X-linked intellectual disability (XLID; Han et al., 2017; Raymond et al., 2007; Schirwani et al., 2018; Masurel-Paulet et al 2014). Individuals with *ZDHHC9* mutations are susceptible to a combined phenotype of speech and language impairments, cognitive difficulties, and Rolandic Epilepsy (RE; Baker et al., 2015). *ZDHHC9* encodes a palmitoylation enzyme involved in the post-translational modification and intracellular trafficking of specific target substrates, including recruitment of receptors and ion channels to the synapse (Fukata & Fukata, 2010). Although multiple substrates may be relevant to neurodevelopmental disorders, one palmitoylation target is thought to be Post-Synaptic Density protein 95 (PSD-95) which is critical to activity-dependent AMPA receptor availability (El-Husseini et al., 2000a; El-Husseini et al., 2000b; Topinka & Bredt, 2010). Palmitoylation is itself activity-dependent, and influences synaptic stability across multiple timescales during development (Kang et al 2008; Kaur et al 2016; Globa and Bamji 2017; Levy and Nicoll 2017). Hence, the loss of *ZDHHC9* function and reduction in palmitoylation efficiency may alter dynamic aspects of post-synaptic activity, impacting on the emergence and stability of functional networks supporting cognition. Because of this proposed physiological role of ZDHHC9, we predicted that mutations to the gene might result in a perturbation to the dynamic nature of large-scale neural networks. Furthermore, we sought to assess whether those networks most altered would reflect the regional expression profile of the gene, whereby the temporal dynamics would be *altered maximally* in those networks in which the gene is highly expressed.

We investigated the dynamics of functional networks at rest and during an auditory oddball task. We included both protocols because it is unclear whether the impact of a mutation upon neuronal dynamics would be most apparent when the system is at ‘rest’ or when under sensory stimulation. For our stimulation protocol we chose an auditory oddball task, because *ZDHHC9* participants have previously been shown to have impaired language development (Baker et al 2015). Auditory oddball tasks, which require rapid habituation to a repeated standard stimulus and sensitivity to deviations from this stimulus, have been used to index auditory processing in developmental language disorders. These studies have established that language impairment is frequently associated with poorer auditory change detection and reduced sensitivity to phonetic and linguistic cues (Ahmmed, Clarke, & Adams, 2008; Baldeweg et al., 1999; Datta et al., 2010; Davids et al., 2011; Shafer et al., 2011).

In sum, the aim of this study was to apply the novel HMM method to map dynamic neural abnormalities in individuals with *ZDHHC9* mutations, during both rest and a passive auditory task, and to determine whether these abnormalities relate to the expression profile of the gene. To our knowledge, this is the first study that has sought to establish whether there is a link between the dynamics of large-scale brain networks, and the regional expression of genes associated with synaptic regulation.

## Methods

### Participants

Eight male participants with an inherited mutation to *ZDHHC9* (age in years: mean = 26.70, standard deviation (SD) = 13.74, range = 13.25-41.83) were compared to seven age-matched male controls (age in years: mean = 27.23, SD = 14.05, range = 10.17-42.50; t = -0.74, p = .943). All control participants had no history of neurological disorders or cognitive impairments. For a detailed description of clinical and cognitive characteristics of the *ZDHHC9* group, compared to an age and IQ matched control group, refer to Baker et al. (2015). In brief, *ZDHHC9* participants had mild to moderate intellectual disability (standardised IQ scores: mean = 64.88, SD = 5.70, range = 57-73). Additionally, they displayed poor verbal fluency, difficulties with non-speech oromotor control, and relatively strong receptive language abilities compared to expressive and written abilities. These communication characteristics distinguished the *ZDHHC9* group from an age- and IQ-matched comparison group (Baker et al., 2015). Ethical approval for the study was granted by the Cambridge Central Research Ethics Committee (11/0330/EE).

### MEG data acquisition and pre-processing

For each participant we acquired data during 9 minutes of eyes-closed resting state and a 12 minute passive auditory oddball task. The data acquired from these two protocols was analysed separately, although using the same pipeline (Figure 1). During the resting-state scan, participants were instructed to relax with their eyes closed and allow their mind to wander, without thinking of anything in particular, but without falling asleep. The oddball task was based upon a design by Cowan et al. (1993) using a roving standard stimulus and delivered in two 6 minute blocks. Participants heard a sequence of tones at the same frequency, followed by a sequence of tones of a different frequency. The repeated tone in each sequence was, therefore, the standard stimulus, and first tone of the new sequence was the deviant tone. The frequency of the repeated standard stimuli altered randomly between three frequencies of 250Hz, 500Hz and 1000Hz. The number of standard stimulus presentations that occurred in a single stimulus train varied randomly from 6 to 12. Tones were 50ms in duration and the inter-tone interval was 500ms. Participants were asked to watch a silent nature documentary and to ignore the tones.

MEG data were acquired at the MRC Cognition and Brain Sciences Unit, Cambridge, U.K. All scans were obtained using the 306-channel high-density whole-head VectorView MEG system (Elekta Neuromag, Helsinki), consisting of 102 magnetometers and 204 orthogonal planar gradiometers, located in a light magnetically shielded room. Data were sampled at 1 kHz and signals slower than 0.01 Hz were not recorded. Before acquisition a 3D digitizer (Fastrack Polhemus Inc., Colchester, VA, USA) was used to record the positions of five head position indicator (HPI) coils and 50–100 additional headshape points evenly distributed over the scalp, all relative to the participants’ fiducial points (nasion and left and right preauricular), which could later be co-registered on MRI scans for source reconstruction. An ECG electrode was attached to each wrist to measure the pulse, and we attached bipolar electrodes to obtain horizontal and vertical electro-oculograms (HEOGs and VEOGs). Head position was monitored throughout the recoding using the HPI coils.

External noise was removed from the MEG data using a signal-space separation (sss) method, and adjustments in head position during the recording were compensated for using MaxMove software, both implemented in MaxFilter version 2.1 (Elekta Neuromag). Following, data were converted to an SPM12 format and down-sampled to 250Hz. Continuous data were inspected and short sections with large signal jumps or artefacts were removed. A temporal independent components analysis (FastICA) was used in sensor space to remove artefacts arising from blinks, saccades and pulse-related cardiac artefacts, by manual visual inspection. The criterion for removing artefacts was a high correlation between the topography of an independent component and one of the HEOG, VEOG and ECG channels. The artefacts were removed by subtraction. Approximately 2-4 components were removed per participant.

**Figure 1.**
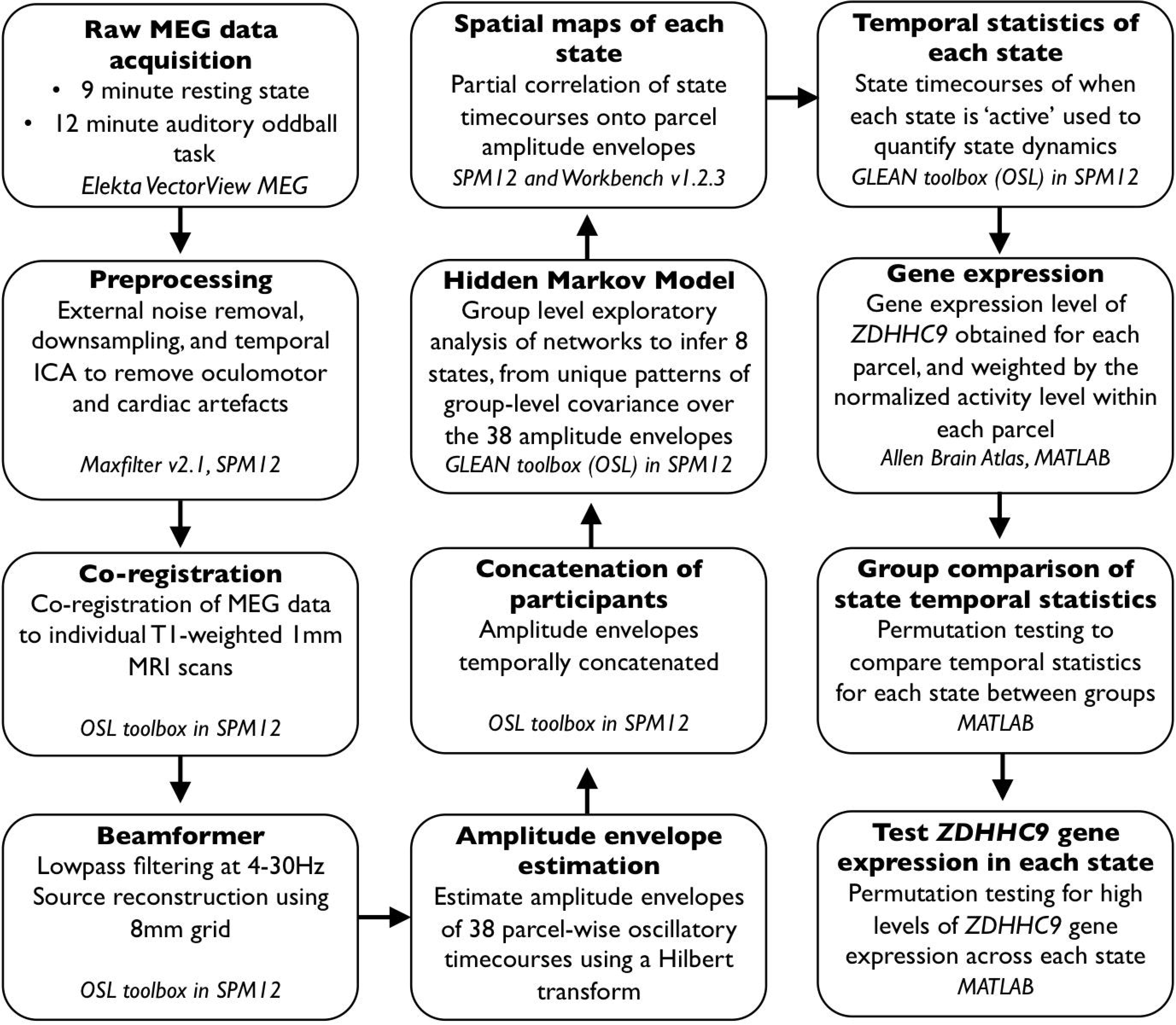
The processing and analysis pipeline used on the resting state and oddball task data.

### MEG source reconstruction, parcellation, and envelope calculation

Each participant’s continuous pre-processed MEG data were co-registered to their individual structural T1-weighted MRI scan with a 1mm image resolution, using the digitised headshape points and fiducials. For one participant without an individual MRI scan, we used a T1-weighted 1mm template in MNI space. Co-registration was run using the OSL toolbox in SPM12 (https://ohba-analysis.github.io/osl-docs/), which aligned the MRI coordinates with the fiducials and headshape points before positioning these data within the MEG sensors using the HPIs for source space analysis.

The MEG sensor data were then low pass filtered at 4-30Hz, to focus on slower frequencies only. This is the frequency range at which amplitude correlations have been shown to produce robust resting state brain networks (Luckhoo et al., 2012; Colclough et al. 2016). For each subject, source space activity was then estimated at every point of an 8mm whole-brain grid using a linearly constrained minimum variance (LCMV) scalar beamformer that combines information across both sensor types and accounts for the reduction in dimensionality induced by the signal-space separation method (Van Veen et al., 1997; Woolrich, Hunt, Groves, & Barnes, 2011). The beamformer used a set of adaptive spatial filters to weight the sensor measurements into an estimate of neuronal activity using the cortical grid, by maximising the band-passed signal at each grid point whilst minimising the signal passed from all other grid points. This process aimed to reduce the effect of signal leakage from neighbouring regions in order to provide a more accurate estimate of activity at each grid location. The beamformer repeated this process across all grid points to produce a whole-brain reconstruction of source-space activity.

Following the beamformer projection of the oscillatory signal into source space, the data across all grid points underwent parcellation into 38 regions of interest using an 8mm mask, applying the method described in Colclough, Brookes, Smith, & Woolrich (2015) and Colclough, et al. (2017). In brief, each parcel was identified using a symmetric orthogonalisation to produce parcel time-courses which were as close as possible to the original time-courses across voxels whilst minimising signal leakage, and correcting for source spread. The time-course of oscillatory activity within each parcel was then obtained using a principal components analysis (PCA), in which the time-courses from all voxels in the parcel were submitted to a PCA, where the first principal component was used as the time-course estimate for that parcel using the spatial filter method.

The amplitude envelope of each parcel’s oscillatory time-course was then calculated using a Hilbert transform, which estimated the instantaneous signal amplitude at each time point. As in Baker et al. (2014), for computational efficiency the amplitude envelopes were down sampled to 40 Hz by temporally averaging within sliding windows with a width of 100ms and 75% overlap between consecutive windows. The parcellation of the source-space data acted to reduce the dimensionality of the oscillatory activity across all source locations into 38 time-courses, to be submitted to the HMM. The amplitude envelopes were then concatenated temporally across all participants to produce a single dataset for the HMM analysis. The envelope data from each participant were demeaned and normalized by their variance prior to concatenation.

### Group level exploratory analysis of networks (Hidden Markov Model)

We ran the HMM using the group-level exploratory analysis of networks (GLEAN) toolbox (https://github.com/OHBA-analysis/GLEAN; Vidaurre et al., 2017). The HMM is specified a priori to derive a certain number of states from the data; we set the HMM to infer eight states, based on previous work suggesting that this number represents a reasonable trade-off between providing a sufficiently rich description and avoiding an overly complex, and therefore hard to interpret, representation (Baker et al., 2014).

The HMM describes the time-series of MEG amplitude envelope data as a temporal sequence of states, where each state describes the data as coming from a unique 38-dimensional multivariate normal distribution, defined by a covariance matrix and a mean vector. Each state, therefore, corresponds to a unique pattern of amplitude envelope activity and covariance that reoccurs at different points in time; where the HMM state time-courses define the points in time at which each state was ‘active’. The estimated state time-courses, corresponding to a binary sequence showing the points in time when that state was most probable, were obtained using the Viterbi algorithm (Rezek & Roberts, 2005).

To produce spatial maps of the changes in amplitude envelope activity associated with each state, the state time-courses were partially correlated onto whole-brain parcel-wise amplitude envelopes concatenated across subjects. The resulting state maps show the brain areas whose amplitude envelopes increase or decrease when the brain visits that state, compared to what happens on average over time.

From the state time-courses, we were able to quantify the temporal characteristics of each state in terms of four measures of interest: 1) Fractional occupancy: the proportion of time each state was active, 2) Number of occurrences: the number of times a state was active, 3) Mean life time: the average time spent in a state before transitioning to another state, and 4) Mean interval length: the average duration between recurring visits to that state. Because these temporal properties reflect the duration and frequency of the coordinated oscillatory activity characterising each state, it was of particular interest to examine whether the expression of the *ZDHHC9* mutation affected the dynamics of these connectivity patterns. To test this hypothesis, we compared each of these temporal characteristics between groups using non-parametric permutation testing. For each temporal property, on each of the eight states, we randomly allocated participants’ values to two groups and calculated the mean difference between these groups. This process was repeated 10,000 times to generate a null distribution of the mean difference between groups that would be expected when group membership was random. The actual group difference was then compared to the permuted null distribution to assess whether this group difference was greater than what we would have expected by chance, and the position of the observed mean difference in the null distribution provided the p-value. All p-values were corrected for multiple comparisons using the Benjamini-Hochberg procedure for false discovery rate (FDR) correction, which is appropriate for maximising power with a large number of tests (Hochberg & Benjamini, 1990).

### ZDHHC9 gene expression permutation testing

Gene expression data were obtained from the Allen Brain Atlas Human Brain public database (http://human.brain-map.org; Hawrylycz et al., 2012). These datasets were based on microarray analysis of postmortem tissue samples from 6 human donors aged between 18 and 68 years with no known history of neuropsychiatric or neurological conditions. MRIs and transformations from individual donor MR space to MNI coordinates were also obtained from the Allen Brain Atlas website. Gene expression (RNA) values were averaged across these 6 donors and mapped onto areas of the Desikan-Killiany (DK) parcellation of the MNI brain. This provided 68 cortical regional *ZDHHC9* expression values (34 in each hemisphere).

Next we wanted to explore whether the impact of the mutation on dynamic transient networks was predicted by the expression profile of the ZDHHC9 gene. Specifically, does regional variation in the expression of the gene predict which networks ought to be most disrupted by the mutation? Simply performing a spatial correlation between the gene expression and the activity pattern across parcels would not work because the networks are highly variable. Instead we calculated a gene expression value for each network, by taking the 20 most active parcels for each network and summing the level of gene expression. The 20 most active parcels of each network were defined as those with the strongest contribution to that network, which was indexed by the absolute partial correlation of the state-time courses with the amplitude envelopes in each parcel. Twenty parcels were selected as this was thought to best reflect an optimal trade-off between peak activation and spatial state distribution.

Before summing them, we weighted the gene expression values by activity levels within each parcel. This weighting was necessary because some networks are heavily dominated by only a few parcels, whereas others are more evenly distributed. To make the process equivalent across all states we normalised the activity levels within each state. The normalisation step allowed us to repeat the process and generate null distributions that were comparable across states. The sum of these values thus provided an expression value for the *ZDHHC9* gene for each network. A null distribution was generated from the random selection of 20 parcel expression values across all 68 cortical parcels, by applying a normalised linear weight activation value, before summing the weighted expression value. This was done for 10,000 iterations to form the null distribution of activation-weighted expression values across each state. We then compared the mean gene expression value of the null distribution compared to the mean for the parcels within each state, to test for a higher level of within-state *ZDHHC9* gene expression than would be expected by chance. Given the use of a smaller number of tests in this case, we corrected for multiple comparisons using the Holm-Bonferroni method, which accounted for the multiple tests done whilst retaining more power than the Bonferroni correction (Abdi, 2010).

## Results

### Dynamic transient networks derived from the HMM in the resting state data

We first ran the HMM on resting data from the control and *ZDHHC9* participants separately, to verify that similar networks were present in each group. The spatial patterns of activity in each state strongly overlapped between groups, and were similar to established resting state networks at slower time-scales (e.g. Damoiseaux et al., 2006; Smith et al., 2012) and to Baker et al (2014).

After confirming that we obtained similar spatial patterns of activity in both groups, we ran the HMM on all participants’ concatenated data to derive common states across both groups, on which we could then compare the groups on temporal measures of interest. Figure 2 shows the spatial maps of each network derived from the resting state data. In the spatial maps, the red/yellow colours represent brain areas in which the amplitude envelope increases when the brain visits that state and blue colours represent brain areas in which the amplitude envelope decreases in that state. The states included sensorimotor networks (State 1 and State 3), a frontoparietal network (State 2), early visual networks (State 4 and State 6), a higher-order visual network (State 5), a left temporal network (State 7), and a distributed frontotemporoparietal network (State 8).

We next characterised the temporal properties of each state in terms of its fractional occupancy, number of occurrences, mean life time, and mean interval length. We tested whether these temporal properties differed between the *ZDHHC9* and control group using non-parametric permutation testing, as described in the Methods, and corrected for multiple comparisons using the Benjamini-Hochberg false discovery rate procedure. On the fractional occupancy measure only an early visual network (State 4) distinguished the two groups, whereby the *ZDHHC9* group spent a higher percentage of time in this state than the control group (actual group mean difference 6.92%; p = 0.01). No other states differed significantly between the groups in fractional occupancy. In terms of the number of occurrences, none of the eight states differed between the groups. The mean lifetime of the states differed between the groups on the early visual network (State 4), where the duration spent in this state was longer for the *ZDHHC9* participants than controls, but this was non-significant following multiple comparison correction. No other states differentiated the groups.

Table 1 presents the descriptive statistics across all states, and Table 2 summarises the group differences and statistical results.

**Figure 2.**
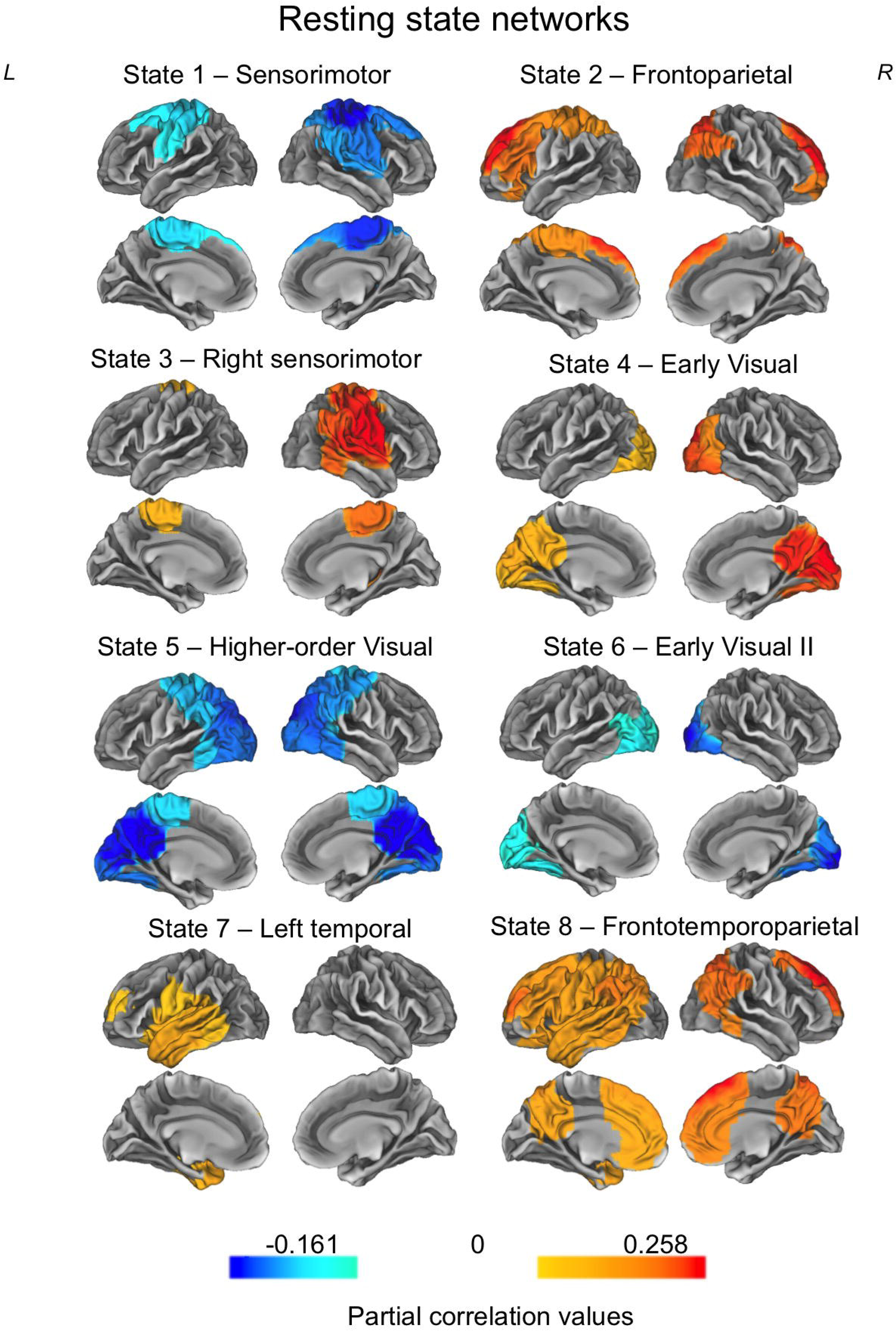
The eight states inferred from the resting state data. Each map shows the partial correlation between each state time course and the parcel-wise amplitude envelopes. The partial correlation values have been thresholded to show correlation values above 70-80% of the maximum correlation for each state, and the colour maps are normalised relative to all states.

**Table 1.**
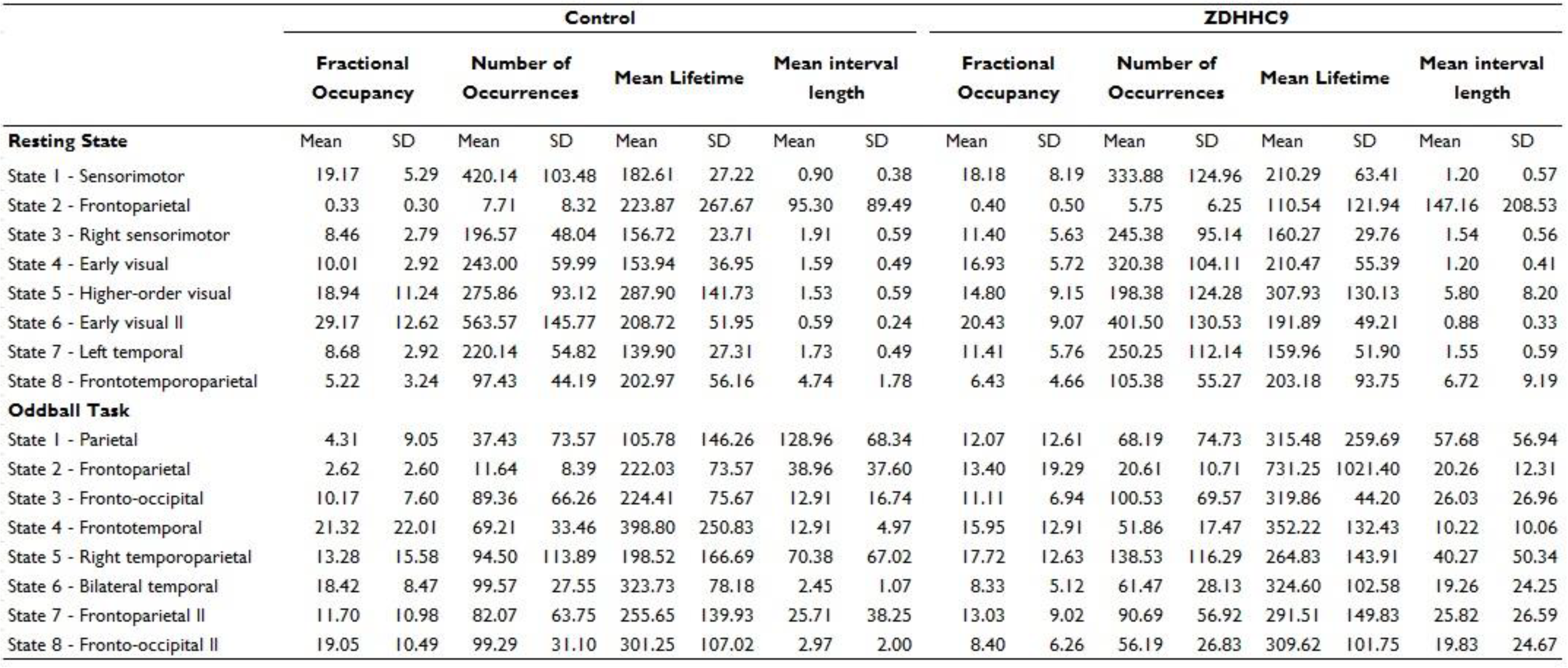
Descriptive statistics of the temporal properties of each network, across the control and *ZDHHC9* participants.

**Table 2.**
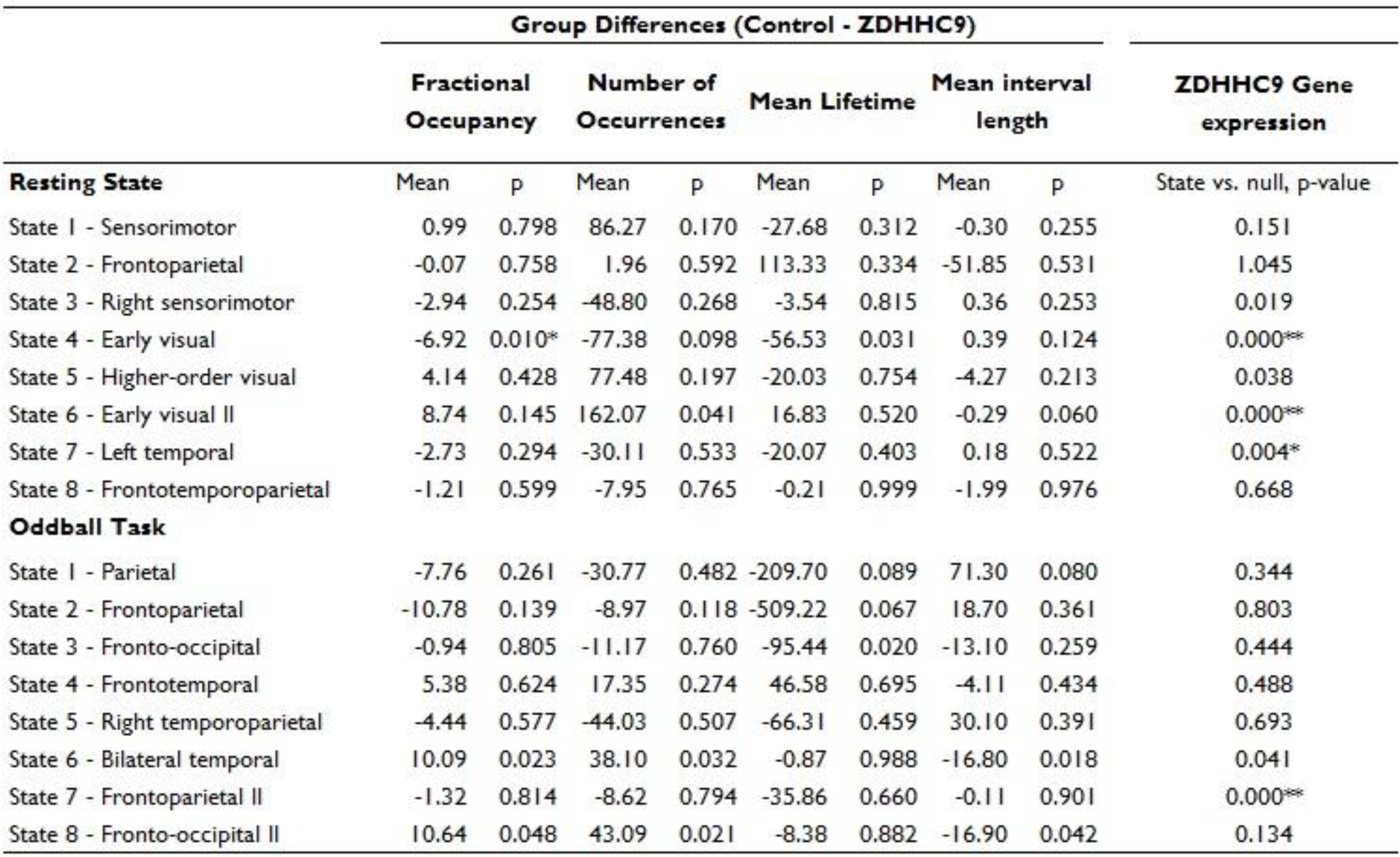
The results of the statistical tests for between-group differences in temporal dynamics across the resting state and oddball tasks, and the statistical test of significant levels of *ZDHHC9* gene expression in each network. Statistical significance was derived using permutation testing and corrected for multiple comparisons as described in the Methods. * denotes significance following multiple comparison correction at the p < .05 level, and ** denotes significance at the p < .001 level.

### Dynamic transient networks derived from the HMM in the oddball data

We applied the same analysis pipeline as described in the Methods to the oddball dataset. Two *ZDHHC9* participants were excluded from the oddball analysis due to failed source reconstruction, but this did not affect the age-matching of the samples (*ZDHHC9* group age in years: mean = 27.11, SD = 12.80, range = 13.25-41.83; comparison with control group, t = -0.015, p = .988). As with the resting state data, we first established whether similar networks were generated by conducting the analysis on each group separately. Following this we combined the two groups into a single analysis. The 8 states derived from the oddball dataset can be seen in Figure 3. The states included a parietal network (State 1), frontoparietal networks (State 2 and State 7), fronto-occipital networks (State 3 and State 8), a frontotemporal network (State 4), right temporoparietal network (State 5), and bilateral temporal network (State 6).

**Figure 3.**
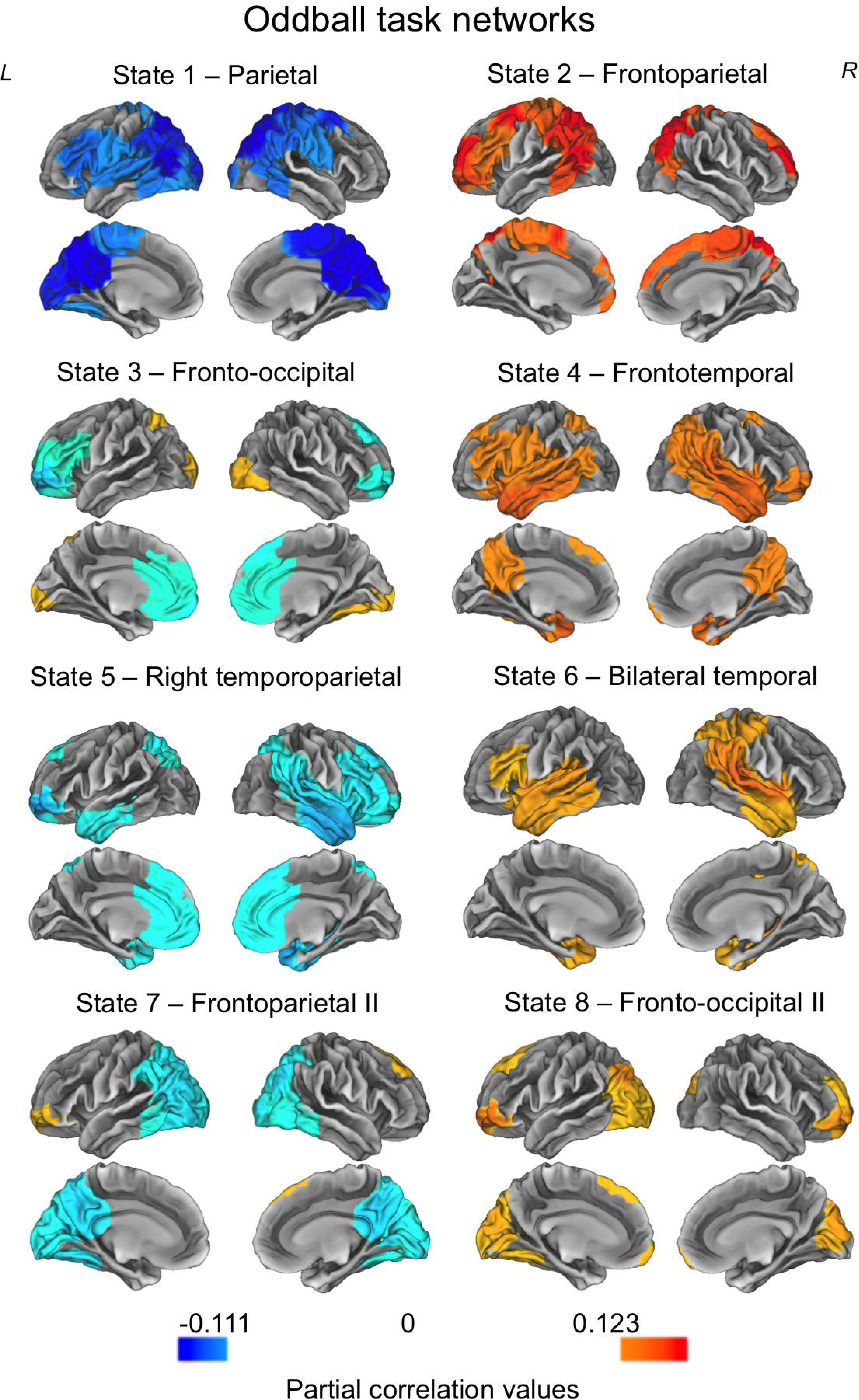
The eight states inferred from the oddball task data. Each map shows the partial correlation between each state time course and the parcel-wise amplitude envelopes. The partial correlation values have been thresholded to show correlations above 60-80% of the maximum correlation for each state. The colour maps are normalised relative to all states.

We next examined whether the temporal properties of the networks distinguished the *ZDHHC9* and control groups. As with the resting state data we tested this using non-parametric permutation testing, as described in the Methods, and corrected for multiple comparisons using the Benjamini-Hochberg false discovery rate procedure. Table 1 summarises the descriptive statistics and Table 2 presents the statistical comparisons between groups. In short, the groups consistently differed in the bilateral temporal network (State 6) and fronto-occipital network (State 8) in fractional occupancy, number of occurrences and mean interval length. The *ZDHHC9* group spent a lower proportion of time in both states, with a lower number of occurrences and shorter mean interval length. However, these group differences were not significant after correcting for multiple comparisons.

In summary, we were able to derive networks from both the resting-state and oddball data which showed good similarity to known functional connectivity networks obtained from fMRI and time-averaged MEG data. The two groups differed significantly in an early visual network in the resting state data, with *ZDHHC9* participants spending a longer period of time in this state. Bilateral temporal and fronto-occipital networks showed descriptive between-group differences, with lower activation indices in the *ZDHHC9* group, but these differences were not significant following multiple comparison correction. The next step in our analysis was to explore the level of *ZDHHC9* gene expression in the HMM-derived networks, and test whether relative between-group differences in network dynamics were associated with the relative expression profile of the gene.

### State-wise gene expression permutation testing

We next wanted to test whether the impact of the mutation on brain dynamics was predicted by the expression profile of the gene. Using gene expression data from the Allen Atlas (Hawrylycz et al., 2012) we identified the level of *ZDHHC9* gene expression within each of the 68 cortical parcels. For each network we could then calculate a gene expression value across the top 20 most active parcels in that state which was weighted by the distribution of activation within each state, as described in the Methods. For each state, the thresholded percentage value refers to the absolute partial correlation threshold required to provide the top 20 active parcels in that state. We corrected for multiple comparisons using the Holm-Bonferroni procedure (see Methods).

**Figure 4.**
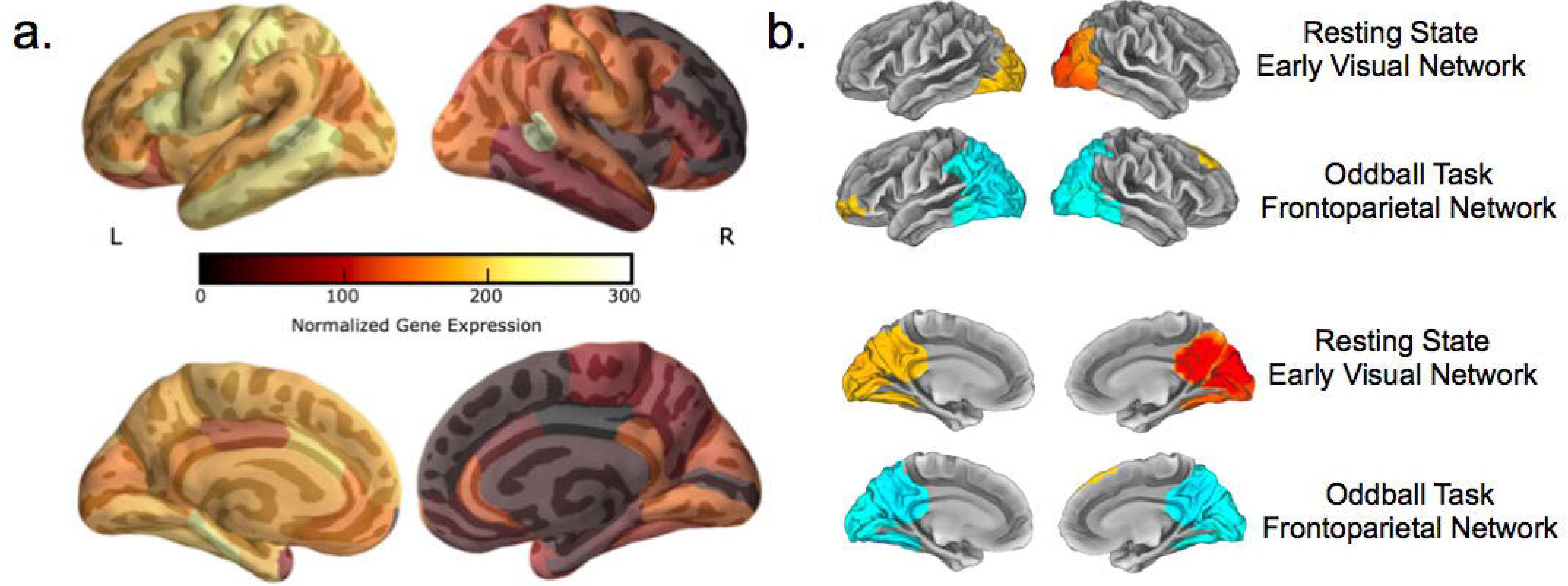
a) The normalized gene expression maps showing the distribution of levels of *ZDHHC9* gene expression throughout the cortex. Lighter colours correspond to higher levels of gene expression. b) Illustrative state maps showing two networks which had significantly above-chance levels of *ZDHHC9* gene expression: the early visual network at rest and the frontoparietal network in the oddball task. The state maps show that the these networks overlap with regions of high gene expression on the normalized gene expression maps.

In the resting state data, the two early visual networks (States 4 and 6) and the left temporal network (State 7) had significantly above-chance levels of *ZDHHC9* gene expression. In the oddball task data, the bilateral temporal network (State 6) and frontoparietal network (State 7) had significantly elevated levels of *ZDHHC9* gene expression. This was relative to what we would expect in a network of that size and activity profile *by chance*, whereby chance levels were derived using permutation testing as described in the Methods. Table 2 presents the results of the statistical tests for gene expression across all states.

In sum, in both the resting state and task data, the gene expression analysis indicated that five of the MEG functional networks identified in both case and control groups showed a disproportionately high expression of *ZDHHC9*: two early visual networks and left temporal network at rest, and the bilateral temporal network and frontoparietal network in the oddball task. Figure 4 presents the cortical distribution of *ZDHHC9* and illustrative state maps showing spatial overlap with regions of high gene expression.

### Statistical association between gene expression and dynamic network group differences

Finally, we sought to determine whether *ZDHHC9* gene expression would predict group differences in neuronal dynamics across networks. We used a Spearman’s rank-order correlation to test for a statistical association between the magnitude of the gene expression effect, indexed by the p-value of gene expression in each state from our permuted distributions, and the magnitude of the group difference, indexed by the p-value of the group difference on each temporal measure (from the group-level permutation testing). Because the p-values were derived from permuted distributions they provided a measure of the *size* of the gene expression effect and group difference in each state, allowing us to test for a relationship between the extent to which network dynamics distinguished the groups and the level of gene expression.

The resting state data showed a significant association between the magnitude of dynamic differences and gene expression. The group difference in fractional occupancy, number of occurrences, and mean interval length was significantly associated with gene expression across states (fractional occupancy: r_s_ = 0.89, p = 0.005; number of occurrences: r_s_ = 0.73, p = 0.048; mean interval length: r_s_ = 0.80, p = 0.022). This demonstrated that these dynamic properties of resting state networks were significantly associated with the strength of gene expression, whereby higher gene expression was associated with a larger impact of the mutation on the dynamic properties across networks. On the other hand, mean lifetime was not significantly associated with gene expression (mean lifetime: r_s_ =0.22, p = 0.606).

The oddball task data did not show a significant relationship between the magnitude of dynamic group differences and gene expression (fractional occupancy: r_s_ = 0.00, p = 1.00; number of occurrences: r_s_ = -0.05, p = 0.935; mean lifetime: r_s_ = -0.55, p = 0.171; mean interval length: r_s_ = 0.24, p = 0.582). Figure 5 presents these results across the resting state and oddball task data.

In sum, we tested the association between group differences in dynamic network properties and gene expression across the resting state and oddball task networks. At rest the size of the group difference in fractional occupancy, number of occurrences, and mean interval length was systematically associated with the level of gene expression.

**Figure 5.**
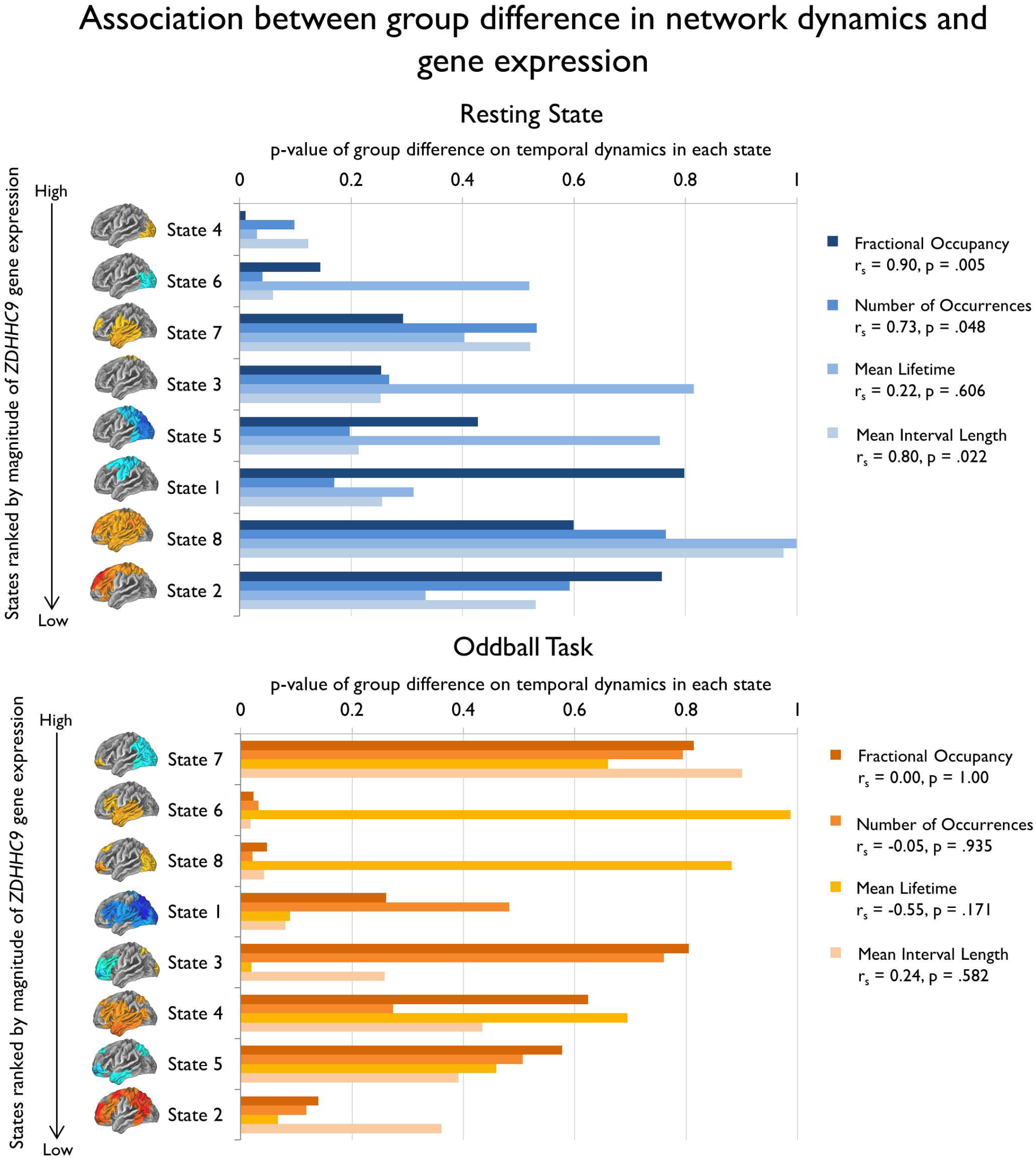
The association between the magnitude of the group difference in network dynamics and the level of *ZDHHC9* gene expression, across all temporal measures and states. The states are ranked by the level of gene expression on the *y*-axis, in which states with smaller p-values derived from the permutation testing have higher levels of gene expression. The *x*-axis shows the p-value of the group difference in network dynamics on each temporal measure, whereby smaller p-values denote larger group differences. The plot for the resting state data demonstrates that the extent to which the dynamic properties differed between groups was associated with a larger magnitude of gene expression across all states. There was no significant across-state association between gene expression and dynamic network group differences in the oddball task.

## Discussion

Little is known about variability in dynamic network properties or the underlying physiological mechanisms that drive this variability. In the current study we sought to characterise the dynamics of functional connectivity networks in individuals with the *ZDHHC9* gene mutation, a single-gene developmental disorder associated with a homogenous phenotype of intellectual, language, and attentional impairments. We examined network dynamics using a Hidden Markov Model (HMM), a data-driven method that captures functional networks on a millisecond timescale and quantifies their dynamics. At rest, an early visual network was active for a significantly longer proportion of time in individuals with the *ZDHHC9* mutation than in age-matched controls. Importantly, across all resting state networks the size of the mutation effect was strongly predicted by the expression profile of the gene: the greater the gene expression within a particular network, the stronger the impact of the mutation on the dynamics of that network. During the auditory oddball task there were no consistent group effects that survived multiple comparisons correction, and there was no correspondence between network dynamics and gene expression. In summary, in resting state MEG data we identified the impact of the gene mutation on the dynamics of specific networks, and crucially, the graded impact of this mutation was predicted by the level of gene expression across networks. However, these gene effects were not reliably detected in task-positive data.

At rest, the higher proportion of time spent in the early visual network in the *ZDHHC9* group suggested less dynamic regulation of transitioning into and out of this brain state than in control participants. One interpretation of this finding is that slower engagement of certain functional networks at rest may present a constraint on higher-level cognitive abilities (Basten et al., 2015). A growing number of studies have argued that the rapid and transient organisation of multiple network configurations at rest necessarily provides the flexibility to adapt to the changing demands of cognitive processing, by providing a continuous ‘dynamic repertoire’ of states to quickly engage the optimal network configuration for a given task (e.g. Bressler & Tognoli, 2006; Deco et al., 2011). Aligning with this, recent work by Schultz and Cole (2016) observed that smaller changes in functional connectivity patterns between rest and distinct tasks correlated with higher levels of fluid intelligence. Given the intellectual difficulties in individuals with the *ZDHHC9* gene mutation (Baker et al., 2015), it is plausible that slower dynamic transitions between resting state networks may constrain the emergence of higher level cognitive abilities through reduced efficiency in coordinating relevant network configurations (e.g. Hearne et al., 2016; Song et al., 2008; van den Heuvel, Stam, Kahn, & Pol, 2009).

However, it is notable that the *ZDHHC9* and control group were only distinguishable on the resting-state dynamics of this early visual network, rather than more distributed frontoparietal or temporal networks which we may have expected to mediate the intellectual and language difficulties in the disorder (e.g. Langeslag et al., 2013). Although functional variations in the occipital cortex have been linked to variability in intelligence (Jung & Haier, 2007) there is weak meta-analytic support for this view (Basten et al., 2015). A plausible interpretation for the present study is that these sensory spatiotemporal patterns of activity were more stable across participants (Lee & Frangou, 2017; Moussa et al., 2012), allowing more reliable measurement of between-group differences in temporal dynamics. The relationship between visual network connectivity and performance across varying language, working memory and reasoning tasks (Schultz & Cole, 2016) suggests that early visual network activation may reflect non-specific engagement in the attentional demands of the study. Nevertheless, the slower dynamics may also be informative about the properties of resting state networks in this disorder.

An important advance in the current study was our demonstration that the integrity of network dynamics strongly overlap with regional differences in *ZDHHC9* gene expression. We predicted that state dynamics would be most altered in regions of elevated gene expression, with these regionally-specific abnormalities potentially arising due to reduced palmitoylation and post-synaptic dysfunction (Fukata & Fukata, 2010; El-Husseini et al., 2010a, 2010b). Our resting state results strongly aligned with this expectation: at rest the magnitude of group differences in dynamic network properties was associated with the level of *ZDHHC9* gene expression across networks, suggesting that higher expression of the *ZDHHC9* mutation has a relatively direct effect on neuronal dynamics. The significantly elevated levels of *ZDHHC9* gene expression in resting state networks, showing case-control differences, suggest that reduced capacity for activity-dependent post-synaptic change, resulting from reduced palmitoylation, may be a contributory mechanism to slower network dynamics (El-Husseini et al., 2010a, 2010b).

There was a striking difference between the resting and oddball data, in that there was no association between *ZDHHC9* gene expression and network dynamics in the oddball task. One possibility is that mutation effects do affect task network dynamics, but that the current sample had insufficient power to detect them. For example, we observed differences in the bilateral temporal and fronto-occipital network dynamics between the control and *ZDHHC9* group. Whilst these effects were not statistically significant, they align with findings in other populations with partially overlapping phenotypes of language impairment who have also shown reduced sensitivity to linguistic and non-linguistic auditory contrasts (Davids et al., 2011), which are in turn predictive of subsequent language development (Port et al., 2016). Furthermore, weaker functional integration in static language-relevant connectivity networks is also a cardinal feature of children with Rolandic epilepsy and comorbid language delays, features observed in the *ZDHHC9* sample (Besseling et al., 2013; McGinnity et al., 2017). The reason why these effects were not robust enough to survive our multiple comparisons correction might be linked to the small number of cases within our sample. However, this would not explain the complete lack of relationship with gene expression in the oddball data, suggesting that this is not simply a power issue. A second alternative is that the measures of neuronal dynamics used here are not the most sensitive for task-positive datasets. There may be dynamic network properties, such as the temporal overlap of activation between states, which we did not detect here but are nevertheless susceptible to disruption from post-synaptic dysfunction. In essence, different neuronal mechanisms could reflect the behavioural phenotype that we observe, but these are not directly influenced by the known aetiology of the *ZDHHC9* mutation.

There are important broader limitations to the current study. First, the sample size of studies of single-gene mutations is inherently limited due to the rarity of these disorders. The present study included all known UK families diagnosed with *ZDHHC9*-associated XLID at the time of recruitment, but the small sample size necessitates replication of these results in a larger sample. However, similar studies of small groups with a common aetiology have generated important hypotheses about neural and genetic pathways to a disorder, such as Fragile X syndrome (Diersson & Ramakers, 2006; van der Molen et al., 2014). The present study further adds dynamic network irregularities as neurobiology-relevant metrics associated with cognitive impairment. Second, the control group was age-matched to the *ZDHHC9* group but had age-typical IQ and language abilities. It is therefore unclear whether the observed dynamic network differences are specifically tied to the combined phenotype of low IQ and language impairment, the aetiology of *ZDHHC9* mutations, or reflect more general correlates of low cognitive ability. It is important for future studies to explore dynamic functional connectivity in individuals with different aetiologies but matched on partially or fully overlapping phenotypes, to draw firmer conclusions about the relationship between causal pathways, dynamic network properties and behavioural phenotypes. In particular, relating dynamic network properties to individual differences in cognition during development may assess whether dynamic network irregularities present a risk factor for specific profiles of cognitive deficits.

Whilst there is a strong tradition of examining the impact of genetic effects on static functional connectivity networks through either heritability (Colclough et al., 2017; Demuru et al., 2017; Fu et al., 2015; Posthuma et al., 2005) or associations between gene expression and amplitude envelope coordination in resting state activity (Gordon et al., 2015; Jamadar et al., 2012; Wang et al., 2015), there remains a scarcity of work examining genetic effects on the fast transient dynamics of functional connectivity networks. Investigating network dynamics is particularly compelling to understand how gene expression perturbs communication within large scale brain networks, and downstream impacts on cognitive abilities, due to the profound impact of genetics on synaptic signalling (e.g. Forest et al., 2017; Meda et al., 2015; Richiardi et al., 2015; Whitaker et al., 2016; Willemsen et al., 2013). For the first time we were able to demonstrate that dynamic connectivity profiles of individuals with a gene mutation are significantly altered and, crucially, that the extent of this alteration is strongly predicted by the expression profile of the gene. Critically, we have demonstrated a valuable method for future work, which should seek to identify the nature of brain network dynamics and their role in the emergence of higher-level cognitive abilities over development.

## Acknowledgements

We thank the study participants, their families, and carers for their extensive contributions and commitment to this project. This study was funded by the Wellcome Trust/Academy of Medical Sciences (Starter Grant for Clinical Lecturers to K. B.). DA and KB are supported by MRC intramural programme grants SUAG/005/RG91365 and SUAG/034/RG91365.

